# RNAseq profiling of leukocyte populations in zebrafish larvae reveals a *cxcl11* chemokine gene as a marker of macrophage polarization during mycobacterial infection

**DOI:** 10.1101/554808

**Authors:** Julien Rougeot, Vincenzo Torraca, Ania Zakrzewska, Zakia Kanwal, Hans J. Jansen, Herman P. Spaink, Annemarie H. Meijer

## Abstract

Macrophages are phagocytic cells from the innate immune system, which forms the first line of host defense against invading pathogens. These highly dynamic immune cells can adopt specific functional phenotypes, with the pro-inflammatory M1 and anti-inflammatory M2 polarization states as the two extremes. Recently, the process of macrophage polarization during inflammation has been visualized by real time imaging in larvae of the zebrafish. This model organism has also become widely used to study macrophage responses to microbial pathogens. To support the increasing use of zebrafish in macrophage biology, we set out to determine the complete transcriptome of zebrafish larval macrophages. We studied the specificity of the macrophage signature compared with other larval immune cells and the macrophage-specific expression changes upon infection. We made use of the well-established *mpeg1*, *mpx*, and *lck* fluorescent reporter lines to sort and sequence the transcriptome of larval macrophages, neutrophils, and lymphoid progenitor cells, respectively. Our results provide a complete dataset of genes expressed in these different immune cell types and highlight their similarities and differences. Major differences between the macrophage and neutrophil signatures were found within the families of proteinases. Furthermore, expression of genes involved in antigen presentation and processing was specifically detected in macrophages, while lymphoid progenitors showed expression of genes involved in macrophage activation. Comparison with datasets of in vitro polarized human macrophages revealed that zebrafish macrophages express a strongly homologous gene set, comprising both M1 and M2 markers. Furthermore, transcriptome analysis of low numbers of macrophages infected by the intracellular pathogen *Mycobacterium marinum* revealed that infected macrophages change their transcriptomic response by downregulation of M2-associated genes and overexpression of specific M1-associated genes. Among the infection-induced genes, a homolog of the human *CXCL11* chemokine gene, *cxcl11aa*, stood out as the most strongly overexpressed M1 marker. Upregulation of *cxcl11aa* in *Mycobacterium*-infected macrophages was found to require the function of Myd88, a critical adaptor molecule in the Toll-like and interleukin 1 receptor pathways that are central to pathogen recognition and activation of the innate immune response. Altogether, our data provide a valuable data mining resource to support infection and inflammation research in the zebrafish model.

## Introduction

Macrophages are phagocytic innate immune cells that, together with neutrophils, form the cellular arm of the first line of host defense against invading pathogens. The activation of macrophages is initiated by the recognition of microbial and danger signals by Pattern Recognition Receptors (PRRs), such as the Toll-like receptors (TLRs), which recruit MYD88 and related adaptor molecules for signal transduction to MAP kinases and Nuclear Factor κB (NFκB) (1). The signaling pathways downstream of TLRs and other PRRs regulate the transcription of a large number of genes that are involved in signaling between immune cells (cytokine and chemokine genes) and in host defense (1; 2). To exert their anti-microbial function, macrophages employ several mechanisms, such as the production of reactive oxygen and nitrogen species, the production of antimicrobial peptides, and the degradation of microbes through the phagosomal-lysosomal and autophagy pathways (3). Following successful elimination of microbial invaders, macrophages mediate the resolution of inflammation by clearing cellular debris and eliminating the surplus immune cells that are undergoing cell death at the infection site (4). In addition to these primary functions in infection and inflammation, macrophages orchestrate a range of developmental processes. For example, macrophages interact with endothelial cells to support angiogenesis (5; 6), help control definitive hematopoiesis (7), and facilitate electrical conduction in the heart (8). Thus, macrophages are highly versatile cells, not only functioning as central players in the immune system, but also contributing to organismal development and maintenance of homeostasis.

Macrophages can adopt different states of activation, which are classically divided into a pro-inflammatory M1 state and an anti-inflammatory M2 state (9). These states are characterized by distinct cytokine and chemokine expression patterns as well as different metabolic profiles. However, it is clear that many intermediate phenotypes exist and that the distinction between M1 and M2 states is a simplification of how macrophage polarization occurs in different organs and tissues and in response to different stress signals (9). Macrophage polarization plays a major role in the context of disease. Tumor-associated macrophages can acquire anti-tumor or tumor-promoting phenotypes (10), macrophage metabolism is critical in development of atherosclerosis and other cardiovascular diseases (11; 12), and certain pathogens are known to manipulate the macrophage phenotype to their advantage (13; 14; 15). Therefore, a better understanding of macrophage polarization and function could open novel therapeutic avenues for diseases related to dysfunction or hyperactivation of this cell type.

The majority of studies on differentiation of macrophage subtypes have been performed *in vitro*, but recently it has been achieved to image the process of macrophage polarization during inflammation in a living organism (16). To this end, the optically transparent early life stages of the zebrafish were used (embryos and larvae), expressing different fluorescent markers for the macrophage cell type and for a classical M1 marker, tumor necrosis factor alpha (*tnfa*). Expression of the *tnfa* reporter was observed in macrophages recruited to sites of injury or sites of *Escherichia coli* infection. Furthermore, the *tnfa*-positive macrophages were shown to express other M1 markers (*il1b*, *il6*), while *tnfa*-negative macrophages expressed M2 markers (*tgfb1*, *ccr2*, *cxcr4b*). Dynamic tracing of reporter expression showed that *tnfa*-positive cells at inflammation sites reverted to a *tnfa*-negative phenotype during the resolution phase (16). In a follow-up study, a tail fin amputation model was used to show that early recruitment of a *tnfa*-expressing macrophage subpopulation is required for blastema formation and subsequent fin regeneration (17). Zebrafish models for a wide variety of human diseases have been developed in the recent years (18). Therefore, the *tnfa*-reporter together with other M1 and M2 lines that are still to be developed, will find many applications to elucidate the functions of polarized macrophage subsets during disease processes.

The most frequently used promoter for driving macrophage-specific expression of fluorescent reporters in zebrafish is that of the macrophage expressed 1 gene (*mpeg1.1*, hereafter called *mpeg1*) (19; 20). The *mpeg1* gene codes for a perforin-like protein with anti-bacterial function (21). Fluorescent *mpeg1* reporter lines have been used to study a diverse range of processes. These include for example, macrophage-endothelial interactions (22), long-distance communication between macrophages and pigment cells (23), the function of tumor-associated macrophages (24), and the role of macrophages in host defense against infections (25). Fluorescent *mpeg1* reporter expression in zebrafish embryos and larvae marks the monocytic precursors of macrophages in the blood as well as tissue-resident macrophages, including microglia in the developing brain (20; 26). The expression of *mpeg1* reporters is non-overlapping with that of a neutrophil-specific BAC reporter line driven by the myeloperoxidase (*mpx*) promoter (27), or with a reporter for immature lymphocytes controlled by the promoter of the LCK proto-oncogene, Src family tyrosine kinase (*lck*) gene (28).

The generation of macrophage and neutrophil reporter lines has gone together with the development of zebrafish models for a variety of human infectious diseases, providing new possibilities to visualize host-pathogen interactions in real time (25; 29). It has been shown that zebrafish embryos rely on macrophages for an effective host defense against different pathogens, such as *Staphylococcus aureus* (30) and *Salmonella enterica* servovar Typhimurium (31). In contrast, the ablation of macrophages was found to protect zebrafish embryos against infection with *Burkholderia cenocepacia*, revealing that macrophages are critical for the proliferation of this pathogen and for the development of a fatal inflammatory response (32). Macrophages play a dual role during infection with *Mycobacterium marinum*, a pathogen widely used to model human tuberculosis in zebrafish, since it provides access to the early stages of tuberculous granuloma formation that is initiated by the aggregation of infected macrophages (33; 34; 35). On the one hand, abundant extracellular growth is observed in macrophage-deficient hosts, indicating that proliferation of *M. marinum* is restricted when phagocytosed by macrophages (36). On the other hand, infected macrophages, driven by bacterial virulence mechanisms, can migrate into tissues and recruit new macrophages, which promotes the cell-to-cell spreading of *M. marinum* and the expansion of granulomas (37; 38). Consequently, a mutation in the macrophage-specific chemokine receptor *cxcr3.2*, which restricts macrophage motility, has a positive outcome on the ability of zebrafish embryos to control *M. marinum* infection (39).

Despite extensive use of the zebrafish *mpeg1* reporter lines for studying macrophage biology, the expression signature of these cells has remained uncharacterized. Here, we isolated *mpeg1* expressing cells from transgenic zebrafish larvae by fluorescence activated cell sorting (FACS) and performed RNAseq to investigate what distinguishes the *mpeg1*-driven expression profile from the signatures of neutrophil and lymphocyte populations isolated from *mpx* and *lck* reporter lines. In addition, we determined a core expression set of 744 zebrafish macrophage markers, based on enriched expression in *mpeg1*-positive cells. We compared this gene set with published RNAseq profiles of human macrophages differentiated in vitro towards M1 or M2 phenotype (40), which showed that zebrafish macrophages express a mixed profile of M1 and M2 markers under unchallenged conditions. We then studied how the expression profile is changed upon *M. marinum* infection and identified a homolog of the human M1 marker CXCL11 as a robust and specific marker of *Mycobacterium*-infected macrophages that is induced by *myd88*-dependent signaling.

## Materials and Methods

### Zebrafish husbandry and infection experiments

Zebrafish lines in this study were handled in compliance with local animal welfare regulations as overseen by the Animal Welfare Body of Leiden University (License number: 10612) and maintained according to standard protocols (zfin.org). All protocols adhered to the international guidelines specified by the EU Animal Protection Directive 2010/63/EU. All experiments with these zebrafish lines were done on larvae before the free-feeding stage. Zebrafish lines included AB/TL, Tg *(mpx:eGFP)*^*i114*^ (27), Tg *(mpeg1:Gal4-VP16)^gl24^*/ *(UAS-E1b:Kaede)*^*s1999t*^ (20), Tg *(mpeg1:mCherry-F)*^*ump2*^ (19), Tg *(lck:eGFP)*^*cz2*^ (28), *cxcr3.2*^−/−*hu6044*^ and their *cxcr3.2*^+/+^ siblings (39), *myd88*^−/− *hu3568*^ and their *myd88*^+/+^ siblings (41). Embryos were grown at 28.5°C in egg water (60 µg/ml Instant Ocean sea salts). *Mycobacterium marinum* M or its RD1-deficient (ΔRD1) isogenic strain (42) containing pSMT3-mCherry (43) was grown and prepared for injections as described (44) and microinjected into the caudal vein of embryos at 28 hours post fertilization (hpf) using, where not differently specified, a dose of 100-125 colony-forming units (cfu) per embryo. After injection, embryos were transferred into fresh egg water and incubated at 28°C for 4 or 5 days before collection. Proper infection was controlled by fluorescent imaging before embryo dissociation.

### Embryo dissociation and Fluorescent Activated Cell Sorting (FACS)

Immune cells from 5-6 dpf larvae were isolated by FACS as described previously (45). Briefly, live embryos were rinsed in calcium free Ringer solution for 15 min and then digested with 0.25% trypsin for 60-90 min at 28°C. Digestion was stopped with 1 mM CaCl2 and 10% fetal calf serum and the cell suspension was centrifuged and washed in PBS. The cell pellet was resuspended in 1 to 2 ml in Leibovitz medium L15 without phenol-red with 1% fetal calf serum, 0.8 mM CaCl2,50 units/ml penicillin and 0.05mg/ml streptomycin and filtered through a 40 µm cell strainer. FACS was performed at 4°C using a FACS AriaIII (BD Biosciences) with the BD FACSDiva software (version 6.1.3). For collecting mCherry-positive cells a Coherent Sapphire solid-state 561 nm yellow green laser with 36 mW power was used. Laser settings applied were 600LP, 615/20 BP. For sorting eGFP and Kaede positive cells a Coherent Sapphire solid-state 488 nm laser with 15.4 mW power was used. Laser settings applied were 505LP, 530/30 BP. mCherry and eGFP/Kaede gates were set-up with non-fluorescent samples and allowed to collect an average of respectively 33.8 +/− 16.4 mCherry and 11.6 +/− 4.4 eGFP/Kaede false-positive cells per million of sorted cells. An average of 526 *mpeg1:Kaede*, 195 *mpx:eGFP* and 983 *lck:eGFP* positive cells in 5 dpf embryos and 1826 *mpeg1:Kaede* and 5482 *mpeg1:mCherry* positive cells at 6 dpf were collected per million of sorted cells. For background expression assessment 500,000 non-fluorescent cells were sorted for each sample. Cell fractions were separately collected in supplemented L15 medium and RNA isolation was performed directly after sample collection.

### RNA isolation, Illumina sequencing, and real time PCRs

RNA extraction was done using the RNAqueous-Micro Kit (Ambion) or RNeasy mini kit (Qiagen). Quality of RNA used for Illumina sequencing was checked on an Agilent Bioanalyzer 2100 using the RNA 6000 Pico kit (Agilent). RNA samples with RIN above 8 were selected. When RNA quantity was low, RNA integrity was judged by the presence of ribosomal peaks. cDNA synthesis and amplification was performed using the SMARTer Ultra Low RNA Kit for Illumina sequencing (Clontech) according to the manufacturer’s protocol. Illumina TruSeq DNA Sample Preparation Kit v2 (Illumina) was used on shared cDNA to prepare libraries. Three changes were made to manufacturer’s protocol: the adapters were diluted 20 times in the adapter ligation step, library size selection was achieved by double Ampure XP purification with a 0.7x beads to library ratio and library amplification was made with 15 cycles. The resulting libraries were sequenced using an Illumina HiSeq2000 with 50 bp paired-end reads for all samples and single-end reads for 6 dpf samples. RNA collected for real time PCR experiments was further purified using column DNA digestion (RNase-Free DNase set, Qiagen). cDNA was prepared using iScript cDNA-synthesis kit (Invitrogen, Life Technologies) and was used as a template for qRT-PCR reaction with iQ SYBR Green Supermix according to the manufacturer’s instructions (Bio-Rad Laboratories). Expression of *cxcl11aa* was analysed using the ΔΔCt method and was normalized against *ppiab* for whole-mount analyses and to *eif5* for FACS sorted cells. Primers for these genes were: *cxcl11aaFw*: ACTCAACATGGTGAAGCCAGTGCT; *cxcl11aaRv*: CTTCAGCGTGGCTATGACTTCCAT; *ppiabFw*: ACACTGAAACACGGAGGCAAAG; *ppiabRv*: CATCCACAACCTTCCCGAACAC; *eif5Fw*: CAAGTTTGTGCTGTGTCCCG; *eif5Rv*: AGCCTTGCAGGAGTTTCCAA.

### RNA-sequencing on 20 sorted cells

Dissociation and cell sorting of infected embryos were performed as mentioned previously. 20 cells were sorted directly in cDNA synthesis buffer from the SMARTer Ultra Low RNA Kit for Illumina sequencing (Clontech) and used directly for cDNA synthesis. The resulting cDNA were amplified for 20 cycles and used for library preparation and single-end Illumina sequencing as mentioned above.

### RNA-seq data analysis

Image analysis and base calling were done using the Illumina HCS version 1.15.1. Quality trimmed reads were aligned to the Ensembl zebrafish genome (Zv9) using Bowtie and reads were mapped to zebrafish transcripts using TopHat and a modified version of the Ensembl Zv9_79 annotation with additional manually annotated genes (Table S1). Differential expression analyses between non-fluorescent cells and fluorescent positive cells were performed using DEseq package in R and DEseq2 for the 20 cell samples. PCA plots and Pearson correlation HeatMaps were generated with DEseq package build in functions. Networks based on GO-enrichment analysis (GOEA) were produced using BiNGO and EnrichmentMap in Cytoscape (46). Gene Set Enrichment Analyses were performed with the GSEA software from the Broad Institute (47; 48) version 3.0.

## Results

### The macrophage-specific transcriptome of zebrafish larvae

To determine the gene expression signature of macrophages, we used *mpeg1*-driven reporters expressing Kaede or mCherry fluorescent proteins (19; 20; 25). RNA was extracted from positive and negative fluorescent cell fractions obtained by FACS sorting of single cell suspensions obtained by dissociating 5 or 6 dpf transgenic larvae. Illumina sequencing (RNAseq) of RNA samples from Kaede-labelled macrophages at 5 dpf and 6 dpf and mCherry-labelled macrophages at 6 dpf, each in duplicate, resulted in a total of 6 replicates. Reproducibility between these replicates after alignment and mapping of the reads was high, as shown by calculation of the Pearson correlation coefficient (Figure S1). Principal component analysis (PCA) showed that all macrophage samples segregated clearly from the samples of negative fluorescent cell fractions. Furthermore, PCA indicated minor differences between samples from macrophages with Kaede and mCherry markers and between the Kaede-labelled macrophages at 5 and 6 dpf (Figure S2A). Between 12110 and 16280 genes were expressed (FPKM ≥ 0.3) in the macrophage populations (Table S2, Table S3) and a total of 13185 genes were shared in at least two of the three conditions. Together, these results indicate that our protocol of RNAseq on FACS sorted cells from zebrafish larvae produces high quality results.

We performed differential expression (DE) analysis on the duplicates from the three different conditions by comparing results from fluorescence positive cell fractions with the related fluorescent negative cell fractions. By selecting an adjusted *p-value* threshold of 0.05, we detected a similar number of genes expressed specifically in Kaede-labelled macrophages at 5 and 6 dpf (715 and 703 genes respectively), whereas more genes were detected in the 6dpf mCherry-labelled macrophages (1953 genes). Comparison between the different conditions showed a high overlap between the enriched gene sets (Figure 1A). To produce a complete and accurate description of the macrophage transcriptome in the larvae, we selected the genes that met the significance threshold in at least two out of the three conditions. This dataset, hereafter named zebrafish macrophage (zfM) core expression set, contains 744 genes (Figure 1A, Table S4).

**Figure 1.**
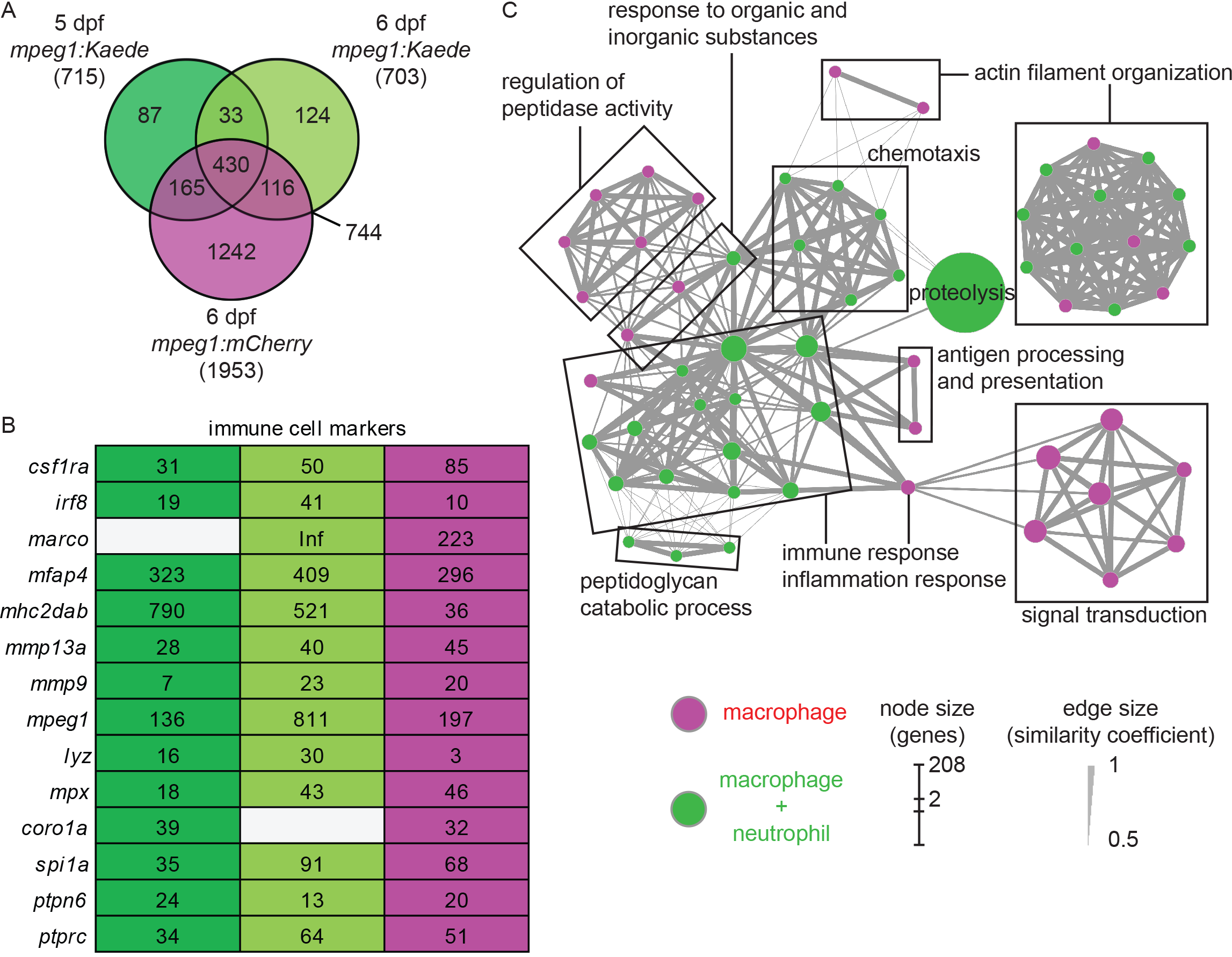
Core expression of macrophages in zebrafish larvae. (A) Overlap of the genes enriched (*P*-adj < 0.05) in zebrafish larval macrophages (zfM) at 5 or 6 dpf from *mpeg1*-driven Kaede and mCherry reporter fish. The zfM core expression is defined as the overlap of two or three *mpeg1* positive populations and includes 744 genes. (B) Expression table of immune cell specific genes. First, second and third columns correspond respectively to macrophage expression from 5dpf *mpeg1:Kaede*, 6dpf *mpeg1:Kaede* and 6dpf *mpeg1:mCherry* reporter larvae. Colored cells correspond to genes enriched in the corresponding sequencing data whereas grey cells correspond to non-enriched genes (log2 (fold change) > 1, *P*-adj ≥ 0.05). Numbers are fold change enrichments in fluorescence positive cells compared to negative cells; Inf = infinite expression change. (C) Network visualization of GO enrichment analysis of genes from the core macrophage expression data set using BiNGO and EnrichmentMap. Red nodes are terms found exclusively in the zfM core dataset and green nodes are found in both zfM core and neutrophil data sets. Node size corresponds to the number of genes associated to the enriched GO term and edge size to the similarity coefficient between two nodes.

The zfM core dataset includes the main genes that are known to be specific for macrophages and myeloid cells in zebrafish larvae (Figure 1B). For example, in addition to *mpeg1* itself, the macrophage-specific genes *csf1ra*, *mhc2dab*, the myeloid genes *spi1a* and *b*, and the pan-leukocyte markers *coro1a*, *ptprc* and *ptpn6* were detected (25; 49). Network visualization of Gene Ontology (GO) terms revealed enrichment of biological processes linked to innate immune response and inflammation, antigen processing and presentation, signal transduction, peptidase activity, chemotaxis and actin filament organization and polymerization (Figure 1C). Similarly, GO terms for molecular function were clearly linked to immune cell function and defense mechanisms (Figure S3).

Among the genes from the zfM dataset, 34% (254 out of 754) corresponded to uncharacterized proteins or non-coding RNAs. Manual annotation showed that many of these uncharacterized sequences belong to large immune-related protein families, including the immunoglobulins (36), the C-type lectins (13) and NACHT/LRR proteins (8) (Table S4). Other uncharacterized genes were also related to immunity, such as genes coding for proteins with chemokine/interleukin-like domains, chemokine receptor like domains, interleukin receptor-like domains, complement domains, leukotriene receptor like domains or MHC class II alpha and beta chains. In addition, 38 genes correspond to non-coding RNAs, long-intronic-non-coding RNAs or processed transcripts, of which the possible role in immunity is of interest for further study.

### Comparison of zebrafish macrophage and neutrophil expression

For comparison with the macrophage transcriptome, we studied the neutrophil transcriptome by sequencing the fluorescent cell population extracted from 5dpf *mpx:gfp* larvae (two replicates). A total of 14241 genes were found expressed in neutrophils (FPKM ≥ 0.3, Table S2, Table S3). Selection of differentially expressed genes revealed a data set composed of 503 neutrophil-enriched genes (*P*-adj < 0.05) (Table S5). 240 (47.71%) of these genes are shared with the zfM core expression dataset (Figure 2A). The neutrophil markers *lyz* and *mpx* were detected enriched at a high level in neutrophil population, although a low level of these transcripts could be detected in zebrafish macrophages (Figure 2A), as well as in human macrophages (40). Similarly, several macrophage markers could be detected in neutrophils, but also at a low expression level (Figure 2A).

**Figure 2.**
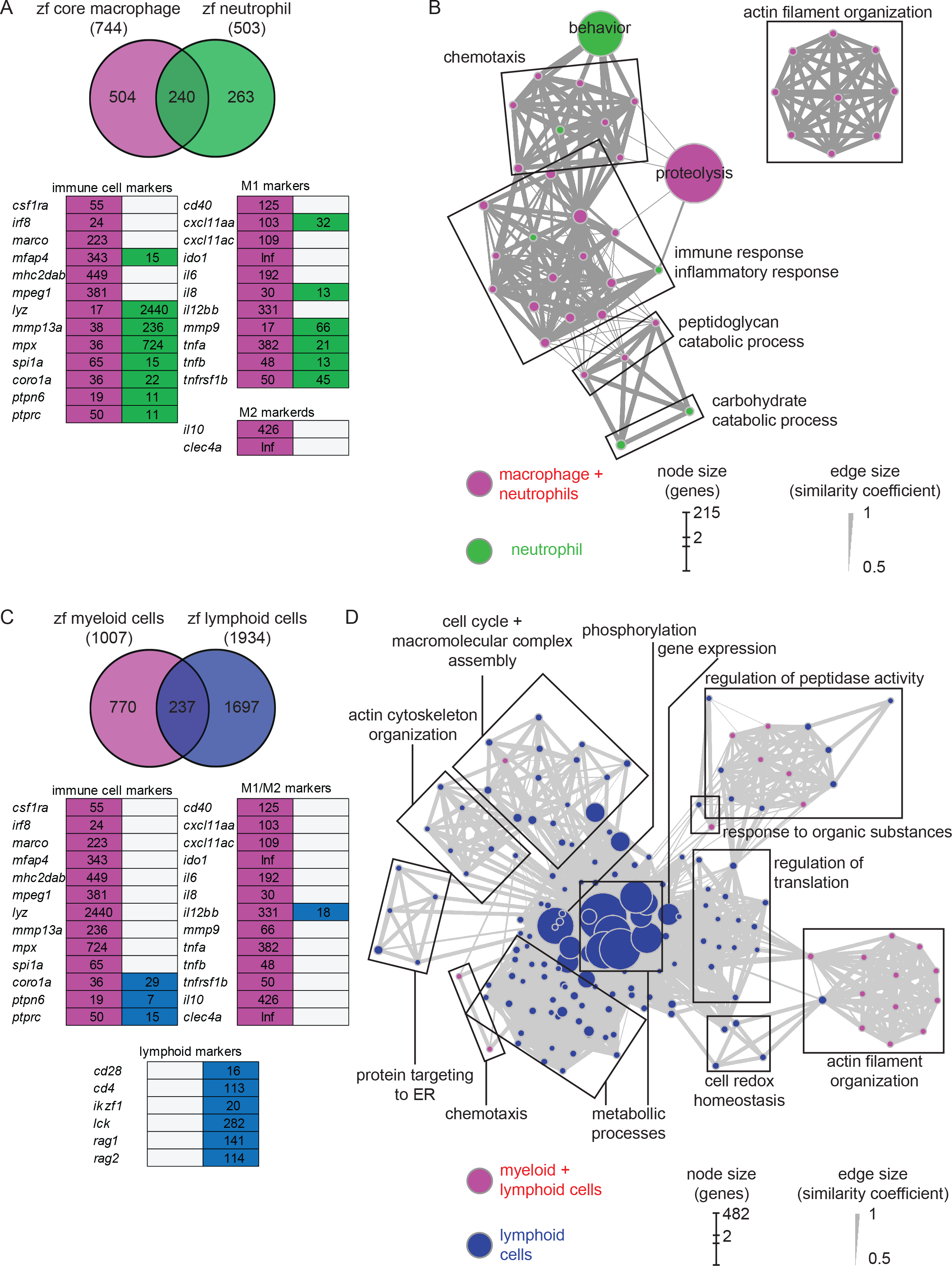
Comparison of macrophage core dataset with neutrophils and lymphoid cells expression sets. (A) Overlap of the genes from the zfM core data set (magenta) and the genes enriched (log2 (fold change) > 1, *P*-adj < 0.05) when comparing *mpx:gfp* positive and negative cells from 5 dpf transgenic fish (green). Below is represented an expression table of a selection of immune cell specific genes. Red cells and green cells represent genes enriched in macrophages or neutrophils, respectively, whereas grey cells represent non-enriched genes. Numbers are fold change enrichments in fluorescence positive cells compared to negative cells; Inf = infinite expression change. (B) Network visualization of GO enrichment analysis of genes from the neutrophil expression data set using BiNGO and EnrichmentMap. Green nodes are terms found exclusively in neutrophil expression dataset and red nodes are found in both core macrophage and neutrophil data sets. Network legend is similar to Figure 1C. (C) Overlap of the genes from the myeloid data set corresponding to genes found either in the core macrophage or the neutrophil expression data sets (red) and the genes enriched (log2 (fold change) > 1, *P-adj* < 0.05) when comparing *lck:gfp* positive and negative cells in 5 dpf *lck:gfp* transgenic fish (blue). Below is represented an expression table of a selection of immune cell specific genes. Red cells and blue cells represent genes detected in myeloid or lymphoid cell populations, respectively, whereas grey cells represent non-enriched genes. Numbers are fold change enrichments in fluorescence positive cells compared to negative cells). (D) Network visualization of GO enrichment analysis of genes from the lymphoid expression data set using BiNGO and EnrichmentMap. Blue nodes are terms found exclusively in lymphoid expression dataset and red nodes are found in both myeloid and lymphoid data sets. Network legend is similar to Figure 1C.

GO analysis identified different biological processes specific for neutrophils such as carbohydrate catabolic process related terms and behavior, described as the internally coordinated responses of whole living organisms to internal or external stimuli and containing the genes *wasb*, *rlbp1b*, *mmp13a* and *cxcr4b* (Figure 2B). GO terms associated to regulation of peptidase activity, signal transduction and antigen processing and presentation are enriched in macrophages but not in neutrophils, confirming the functional differences between the two myeloid lineages (Figure 1B). Many GO terms are shared between the two cell populations. However, their contents often are composed of different protein families (Figure 3). For example, proteolysis appears to be a major group in both cellular lineages, but macrophages express cathepsin coding genes (*ctsba*, *ctsc*, *ctsh*, *ctsk*) whereas neutrophils express proteinases from the carboxypeptidase (*cpa1*, *4*, 5), the elastase (*ela2*, *ela2l* and *ela3l*), the chymase families as well as trypsin.

**Figure 3.**
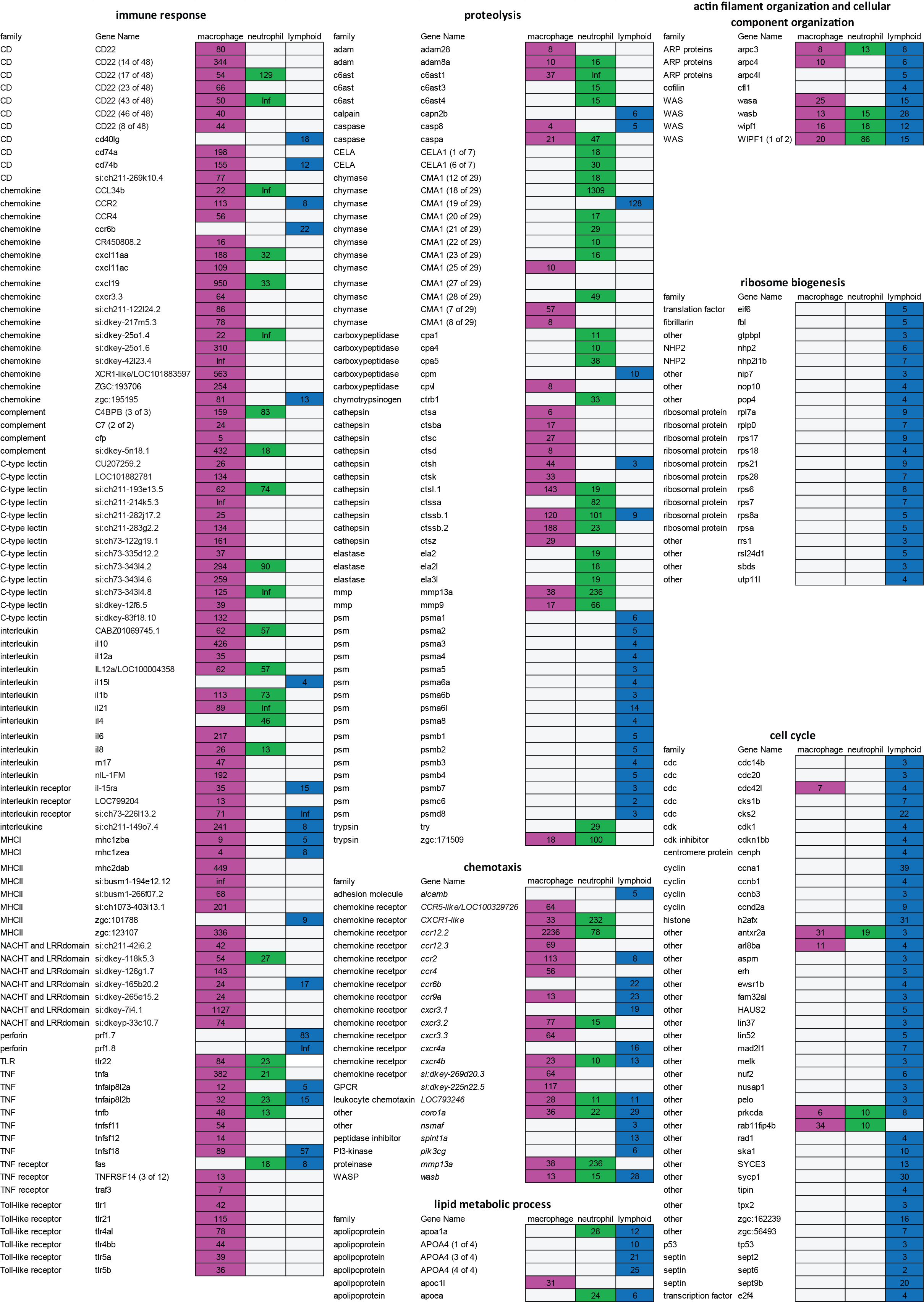
Different gene sets expressed in macrophage, neutrophil and lymphoid cell population. Expression table of selected genes belonging to different GO terms. First, second and third columns correspond to macrophage, neutrophil and lymphoid cell populations, respectively. Magenta, green and blue cells represent genes enriched in macrophages, neutrophils, or lymphoid cells. Numbers are fold change enrichments in the fluorescence positive cell fractions compared to negative cell fractions. Grey cells are non-enriched genes (*P*-adj > 0.05).

### Comparison of zebrafish myeloid and progenitor lymphoid expression

We then used the *lck:GFP* transgenic line to study the lymphoid cell transcriptome. At 5dpf, the adaptive immune system is not yet mature. However, lymphoid cell progenitors are already present in the thymus (50). RNAseq analysis showed that 13153 genes are expressed in this cell population (FPKM ≥ 0.3, TableS2, Table S3) and 1934 genes were enriched in the lymphoid cell progenitor compared to non-fluorescent cell background (Table S6). Comparison of genes enriched in lymphoid cells with genes specific for myeloid cells (i.e. genes enriched in either macrophages or neutrophils) showed a small overlap between the two populations (Figure 2C). None of the macrophage or neutrophil markers was detected in the lymphoid cell transcriptome, except for *il12bb*, although it is less expressed. On the opposite, several known lymphocyte markers were detected only in this cell population (Figure 2C). GO analysis showed also little overlap with processes detected in myeloid cells (Figure 2D). Surprisingly, very few terms were associated to immune function, except for chemotaxis, regulation of peptidase activity and actin filament organization, also shared with the myeloid lineage. However, these groups are composed of different genes. For example, the proteases expressed by lymphoid cells mainly belong to the proteasome (Figure 3). A closer look to immunity-related genes showed the presence of chemokines and chemokine receptors as well as MHC class II genes, but expressed at a low level compared to myeloid cells (Figure 3). Two perforin genes were also highly expressed in lymphoid cells only. The main enriched GO terms in the lymphoid cells were found to be associated with metabolic processes, cell cycle, and ribosome biogenesis (Figure 2D), which might reflect the immature status of this cell population in developing zebrafish larvae.

### Similarity between the zebrafish macrophage transcriptome and human polarized macrophage transcriptomes

By real time imaging of macrophages in a dual fluorescent *mpeg1* and *tnfa* reporter line evidence has been obtained that zebrafish larvae differentiate M1 and M2 like polarized macrophages in response to wounding and infection (16). We found that the known M1 (*il1b*, *tnfa/tnfb*, *il6*) and M2 (*cxcr4b*, *il10*, *ccr2*) markers were expressed in the zfM core expression set. Additionally, the zebrafish homologs of human M1-markers *CXCL11* (*cxcl11aa*)*, IDO1* (*ido1*)*, MMP9* (*mmp9*) *and TNFRSF1B* (*tnfrsf1b*) *and the M2-markers ALOX5AP* (*alox5ap*)*, CLEC4A* (*si:ch211-170d8.8*)*, MARCO* (*marco*) and *TGFB1* (*tgfb1b*) were also detected in our zfM core dataset (Table S4).

To investigate further the similarities with human macrophages, we compared our transcriptomic data with RNA sequencing data published by Beyer et al. (2012) (40), in which transcriptomes of *in vitro* polarized M1 and M2 macrophages were analysed.

By using Biomart (http://www.ensembl.org/biomart/) combined with custom annotations, we retrieved the human homologs of the total of 13185 genes expressed in *mpeg1*-positive macrophages and identified 9780 human genes with a HGNC symbol. Approximately three quarters of this human homolog set were also found among the set of 12327 genes expressed in human M1 cells (73,1%) and among the 12488 genes expressed in M2 cells (73,6%) (RPKM > 0,3) (Figure 4A). Similar proportions of the human homologs from the zfM core expression set (for which 524 human homologs were identified, see Table S7) were found in human M1 (75,8%) and M2 (76,2%) macrophages (Figure 4A). Among those genes, only 11 were expressed exclusively in human M1 macrophages and 13 exclusively in M2 macrophages. These observations suggest that zebrafish macrophages are composed by a mixed population of M1 and M2 type macrophages.

**Figure 4.**
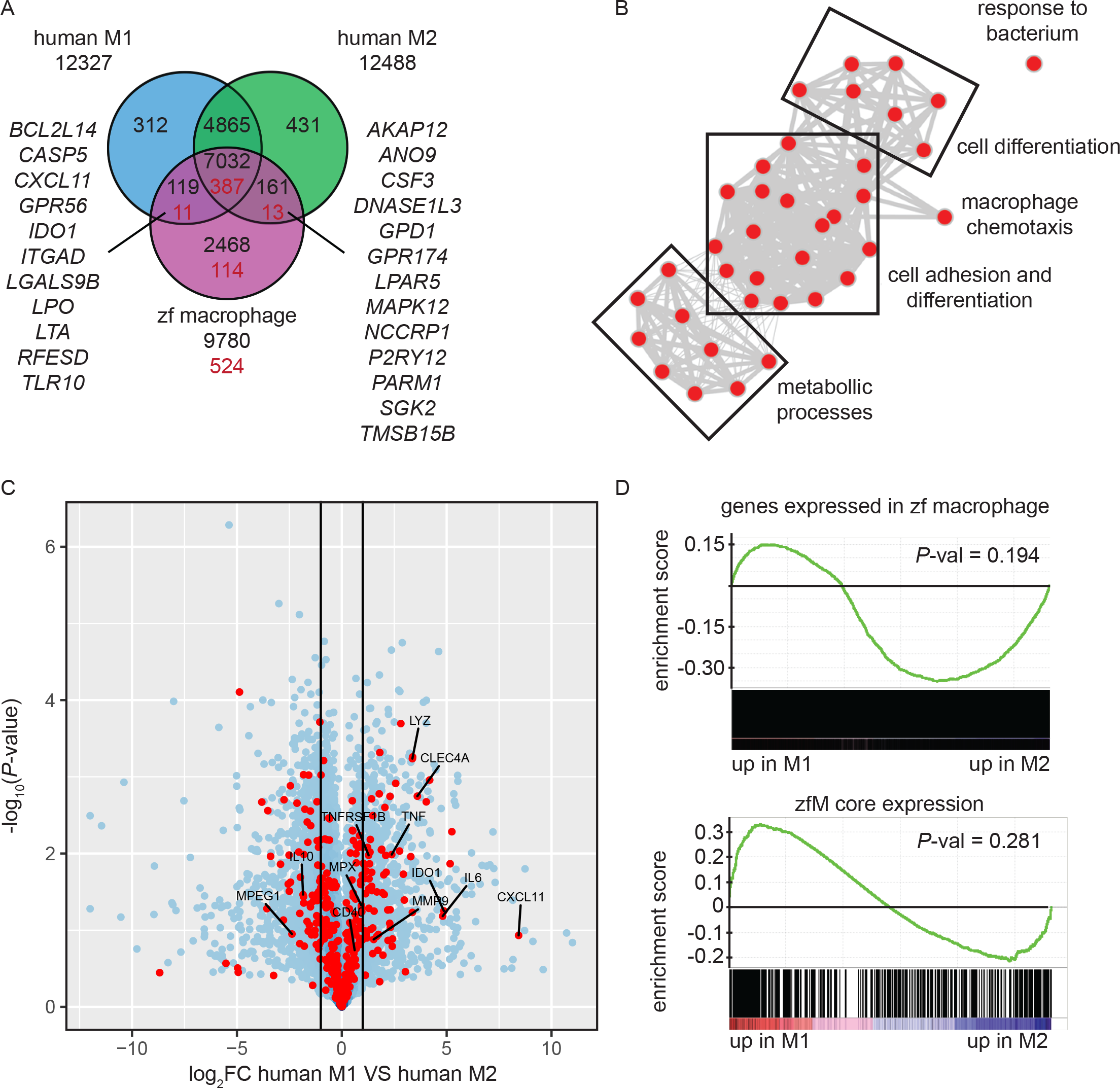
Zebrafish larval macrophages have a gene signature similar to human M1 and M2 polarized macrophages. (A) Overlap of the genes detected in human M1 (blue) or M2 (green) polarized macrophages and of the human homologs of the zebrafish core macrophage data set (magenta). Black number correspond to the comparison between genes expressed in human or zebrafish cells (RPKM or FPKM>0.3) and red numbers correspond to the comparison of gene expressed in human cells and specifically enriched in zebrafish macrophages (log2 (fold change) ≥ 1, P*-adj* < 0.05). (B) Network visualization of GO enrichment analysis of human homologs of zebrafish macrophage enriched genes not detected in the dataset of human M1 and M2 in vitro polarized macrophages using BiNGO and EnrichmentMap. Red nodes represent GO terms. Network legend is similar to Figure 1C. (C) Volcano plot showing the P-value (-log10-transformed) as a function of the fold-change (log2-transformed) between human M1 and M2 gene expression level of the gene set from Beyer et al. (2012). Red dots are genes with a human homolog detected in the zebrafish macrophage core dataset. (D) Gene Set Enrichment Analysis (GSEA) plots of gene expression changes in human M1 *in vitro* polarized macrophages compared to human M2 *in vitro* polarized macrophages from Beyer et al. (2012). Gene sets used for the analyses are genes expressed in zebrafish macrophages (FPKM > 0.3) (top panel) and genes from the zebrafish macrophage core dataset (lower panel).

A total of 114 homologs from the zfM core expression dataset were not present in the M1 or M2 polarized human macrophage datasets (Figure 4A). Among these were the known M1 marker Interleukin 12B and the M2 marker mannose receptor C type 1 (MRC1). Other genes detected exclusively in the zfM expression set were associated to the molecular function carbohydrate binding, serine-type endopeptidase inhibitor activity and NAD(P)+ protein arginine ADP-ribosyltransferase activity, and involved in peptidoglycan catabolic processes (Figure 4B). Genes coding for several members of the SerpinB serine protease inhibitor family, the peptidoglycan recognition protein Pglyrp family, metalloproteases (Mmp13a), as well as cytokines (Tnfs11, Il21) and cytokine receptors (Ccr9a, Cxcr3.2, Cxcr3.3, Il22ra2) are present in these categories. The absence of these genes in the human M1 and M2 sets might be due to low expression levels in *in vitro* cultured cells.

Finally, we computed the differential expression between human M1 and M2 polarized macrophages and searched whether the genes from the zfM core expression set were preferentially associated to either M1 (log2FC > 1) or M2 (log2FC < −1) signal. The results show that 67 genes from the zfM core dataset were associated to M1-enriched genes and 67 were associated to M2-enriched genes (Figure 4C).

We also used Gene Set Enrichment Analysis (GSEA) to compare the set of genes expressed (RPKM > 0.3) in zebrafish macrophages with the differential expression between human M1 and M2 polarized macrophages. The analysis showed a preference for M2-enriched genes, although this enrichment was not significant (Figure 4D top panel). Focusing on the zfM core gene set also showed no clear enrichment for either M1- or M2-enriched genes (Figure 4D lower panel).

Altogether, our results indicate no clear polarization of the zebrafish macrophages, suggesting the presence of both M1 and M2-typed macrophages in unchallenged larvae.

### Effect of *M. marinum* infection on the zebrafish macrophage transcriptome profile

As zebrafish larval macrophages display mixed M1 and M2 characteristics, we tried to induce a shift in activation phenotype by infecting *mpeg1:mCherry* embryos at 1dpf with GFP-labelled *Mycobacterium marinum* (*Mm*), an intracellular pathogen of macrophages. Transcriptomes of infected and uninfected macrophages were profiled 5 day post infection (6 dpf).

When retrieving samples from infected larvae, only a small number of double positive cells were collected over a long sorting period, inducing variation between replicates. To minimize the differences, we reduced the sorting time and the number of steps in the protocol by collecting 20 infected and uninfected cells from Mm-infected larvae directly in cDNA synthesis buffer and by proceeding immediately to cDNA synthesis and amplification without RNA extraction. These modifications of the protocol led to reproducible results (Figure S2, Figure S3). Differential expression analysis between infected and uninfected macrophages identified 330 upregulated and 139 downregulated genes (*P*-adj < 0.05) (Figure 5A, Table S8). GO analysis identified two terms enriched in the downregulated genes only: cell cycle and blood vessel morphogenesis. This group, often related to immune cell migration, included the genes *flt1/VEGFR1*, known to be expressed in mouse M2 macrophages *in vitro* (51; 52), and *ptprja/CD148*, expressed by human macrophages under exposure to LPS and other TLR-ligands but repressed under CSF-1 treatment (53). Performing GO analysis on the human homologs of this set of genes identified the terms transcription coactivator activity, NADP or NADPH binding and serine hydrolase activity associated to upregulated genes (Figure 5B) and protein localization to downregulated genes.

**Figure 5.**
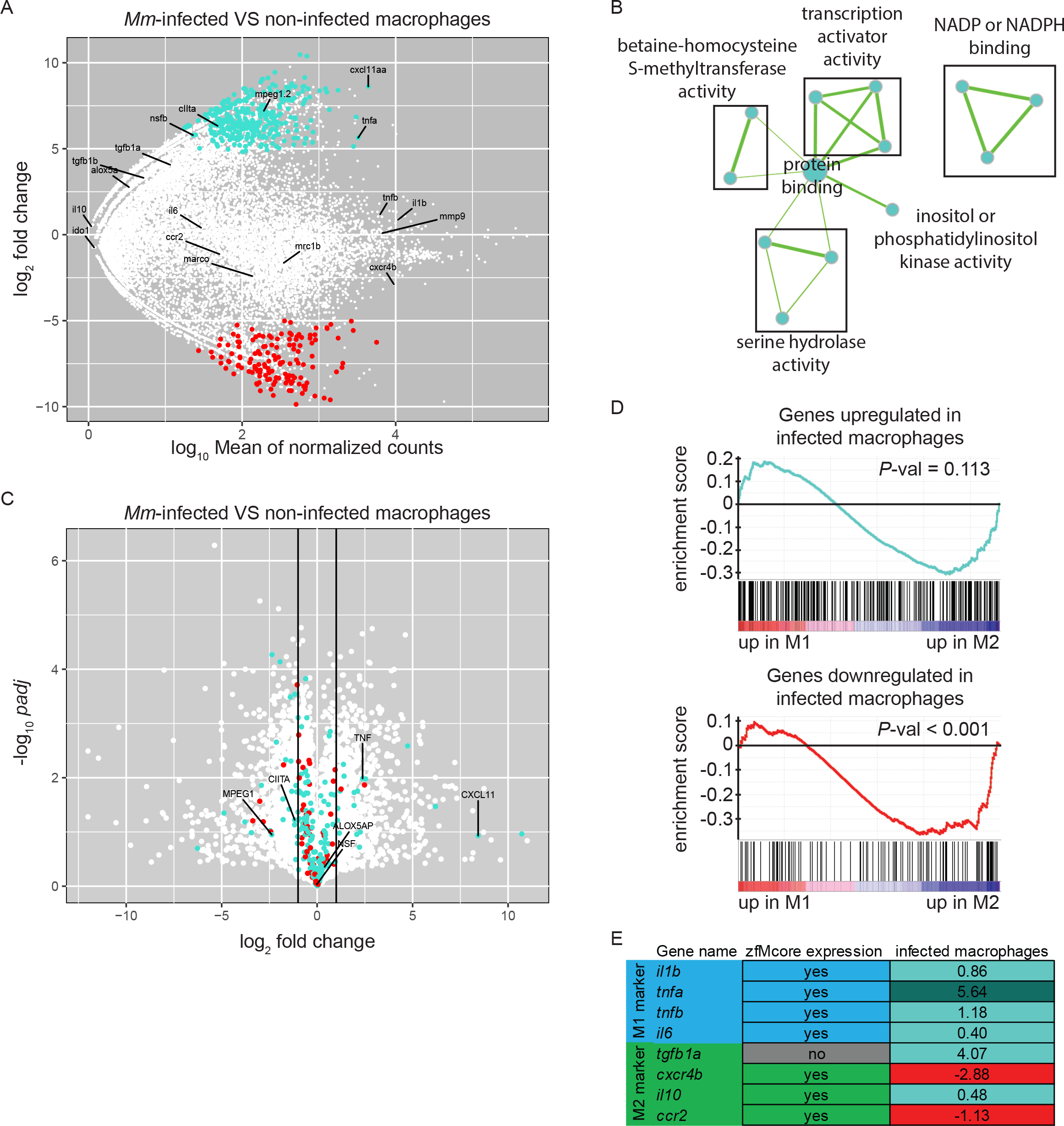
*M. marinum* infected macrophages exhibit a change in gene expression towards M1-polarization. (A) MA-plot showing the fold change (log2-transformed) between gene expression in Mm-infected and non-infected *mpeg1:gfp* positive cells from a 6 dpf embryos 5 days after injection of 100-125 cfu of *Mycobacterium marinum* M strain containing pSMT3-mCherry as a function of the normalized average count between the two conditions (log10-transformed) as calculated with DEseq2. Turquoise: log2FC ≥ 1 and *P*-adj < 0.05, red: log2FC ≤ −1 and *P*-adj < 0.05. (B) Network visualization of GO enrichment analysis of human homologs of up-regulated genes in infected macrophages compared with uninfected macrophages from infected larvae using BiNGO and EnrichmentMap. Network legend is similar to figure 1C. (C) Volcano plot showing the P-value (-log10-transformed) as a function of the fold-change (log2-transformed) between human M1 and M2 gene expression level of the gene set from Beyer et al. (2012). Turquoise and Red dots are genes with a zebrafish homolog respectively up-and down-regulated in the infected macrophages compared with the non-infected macrophages from *M. marinum* infected larvae. (D) Gene Set Enrichment Analysis (GSEA) plots of gene expression changes in human M1 *in vitro* polarized macrophages compared to human M2 *in vitro* polarized macrophages from Beyer et al. (2012). Gene sets used for the analyses are human homologs of genes found up-regulated (log2FC ≥ 1 and *P*-adj < 0.05) (top panel) and down-regulated (log2FC ≤ −1 and *P*-adj < 0.05) (lower panel) in macrophages upon Mm infection as described in (A). (E) Table presenting the zebrafish genes expressed in M1 and M2 macrophages studied in Nguyen-Chi et al. (2015). First column indicates their presence in our zfM core dataset. Second column indicates their enrichment (log2Fold Change) in Mm-infected macrophages compared to uninfected macrophages. Turquoise: log2FC ≥ 1 and *P*-adj < 0.05, light turquoise: log2FC ≥ 0, red: log2FC ≤ 0.

Our analysis revealed several *Mm*-induced genes that could play important roles in host defense. These include for example CIITA, the master transactivator of MHC class II gene expression, which has previously been described to be important for limiting *M. tuberculosis* infection in mice (54). Another Mm-induced gene is the *mpeg1*-family gene *mpeg1.2*, which we have previously shown also to be inducible by *Salmonella* infection (21). The *mpeg1* genes encode proteins of the perforin family with proposed anti-bacterial functions in macrophages that require further mechanistic dissection (21). On the other hand, other overexpressed genes could be more beneficial for the survival of the bacteria. The gene *nsfb*, the zebrafish homolog of the human N-ethylmaleimide sensitive factor, has been proposed to promote the fusion of phagosomes containing live *Salmonella* with the early endosome and repress their transport to lysosomes (55), whereas the *acap1* gene promotes *Salmonella* invasion (56).

To explore the possible polarization of the *Mm*-infected zebrafish macrophages, we compared the differentially expressed gene set with the transcriptomes of M1 and M2 *in vitro* polarized macrophages reported by Beyer *et al.* (2012) (40). We found that 18 M1-enriched genes (log2FC > 1) were overexpressed in infected macrophages and 2 were downregulated whereas 26 M2-enriched genes (log2FC < −1) were upregulated and 6 were downregulated (Figure 5C). Although GSEA showed no significant association of the upregulated genes with either M1- or M2-enriched genes (Figure 5D, top panel), the genes downregulated in *Mm*-infected macrophages were clearly associated to M2-polarized human macrophages (Figure 5D, lower panel).

One of the most highly induced gene in infected macrophages was *cxcl11aa*, a zebrafish homolog of the gene for human CXCL11 (Figure 5A), a proinflammatory chemokine that is a typical M1 marker (57). We recently showed that this chemokine is important during *Mm* infection in zebrafish for the recruitment of macrophages and dissemination of the bacteria(39). Furthermore, expression of *tnfa* appeared to be highly upregulated in infected macrophages. Tnfa is one of the main markers of M1 activated macrophages in human and has been used as a marker for M1-like activated macrophages in zebrafish larvae (16). Other known zebrafish M1-like activated macrophage markers are non-significantly overexpressed (*il1b*, *tnfb*), or barely expressed (*il6*). On the other hand, the known zebrafish M2-like markers are either expressed at a low level (*tgfb1a*, *il10*) or not significantly downregulated (*cxcr4b*, *ccr2*) (Figure 5A,E).

We can conclude that the strong induction of two important proinflammatory markers, *cxcl11aa* and *tnfa*, and the downregulation of genes associated to M2 polarization as detected by GSEA indicate that *Mm*-infected macrophages display M1 rather than M2 characteristics.

### The *cxcl11aa* gene expression as a robust marker of *Mycobacterium*-infected macrophages

Among the chemokine and cytokine genes expressed in *Mm*-infected macrophages, *cxcl11aa* emerged as the most reproducible infection marker from the RNAseq analysis, showing significantly higher induction (average log2 (fold change) = 8.6, *P-*adj < 0.001) than *tnfa* (average log2 (FC) = 5.6, *P*-adj = 0.03) in all replicates. In order to confirm the *Mm*-inducible expression of *cxcl11aa* in macrophages, we FACS-sorted *mpeg1:mCherry* positive cells from *Mm*-infected and mock-injected larvae and quantified the level of *cxcl11aa* expression by real time PCR. In uninfected conditions, the expression of *cxcl11aa* was significantly enriched in the mCherry-positive macrophage cell fraction compared with the unlabeled cell fraction (Figure 6A). During infection, the expression levels of *cxcl11aa* were strongly upregulated in macrophages but not in the unlabeled cell fraction. We found that the level of this infection-induced and macrophage-specific expression of *cxcl11aa* is high enough to be detectable in total RNA samples from whole larvae and that *cxcl11aa* induction did not require the bacterial locus RD1 (Region of Difference 1), a pathogenicity locus encompassing the secretion system of ESAT-6 (Early Secreted Antigenic Target 6 kDa), which is associated with mycobacterial virulence and formation of tubercular granulomas (Figure 6B) (58). The induction of *cxcl11aa* was also independent from the host *cxcr3.2* gene, which encodes the receptor for Cxcl11aa (Fig.6B) (39). Next, we asked whether *cxcl11aa* induction requires the central immune mediator Myd88, which links pathogen recognition by Toll-like receptors and Il1β-mediated inflammation to activation of the transcription factor Nfκb (59). Therefore, we quantified the expression levels of *cxcl11aa* in *myd88* mutant larvae. Since *myd88* mutants display an increased infection level when infected with the same initial infection load as wild type siblings, we compensated this with a reduced inoculum to obtain a similar infection level at 4 dpi. Both with the reduced and the regular inoculum, *myd88* mutants displayed a marked incapability to upregulate *cxcl11aa* (Figure 6C-E), indicating that Myd88-dependent signaling is key to upregulate macrophage expression of *cxcl11aa* during *Mm* infection.

**Figure 6.**
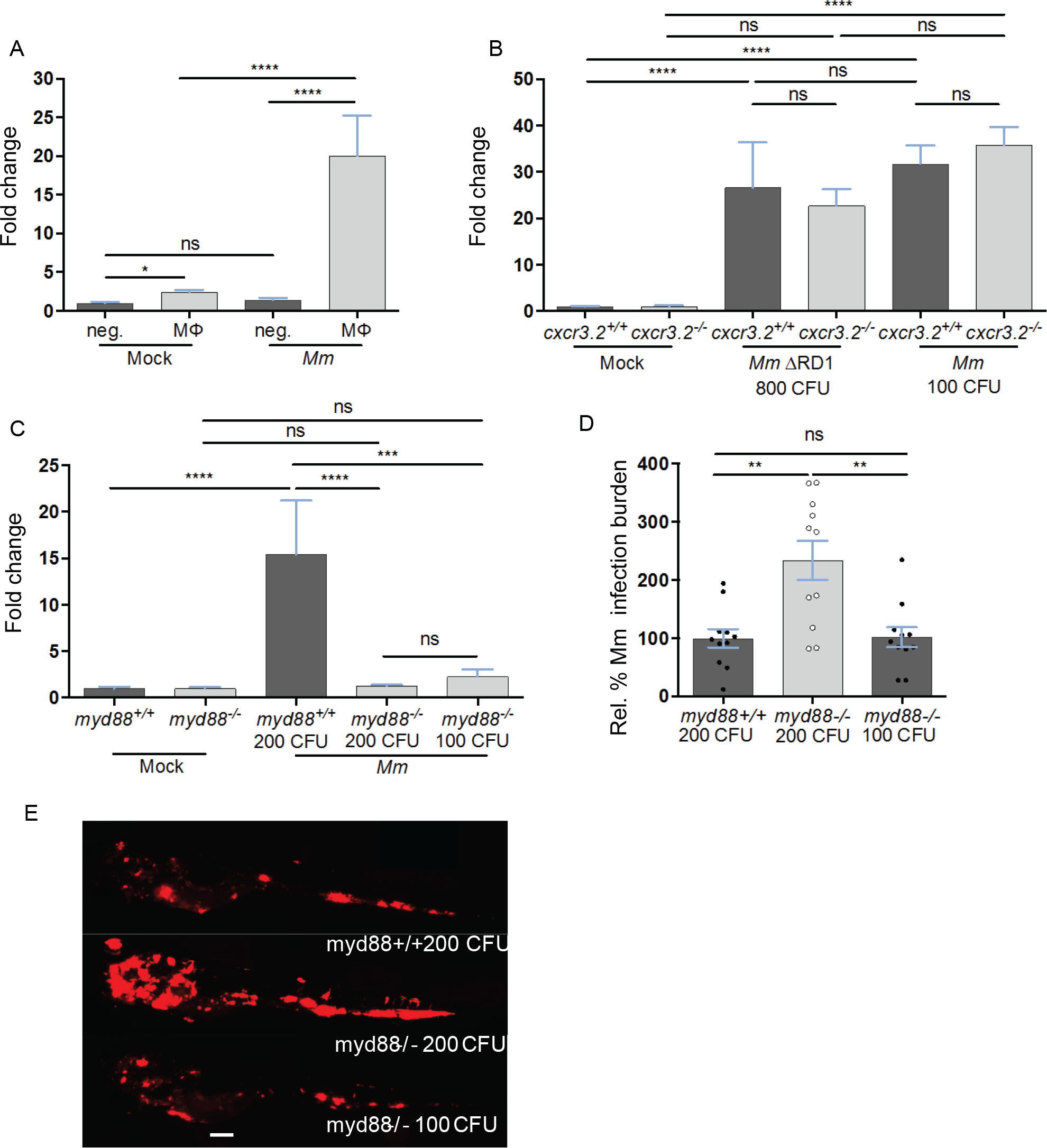
The expression of *cxcl11aa* is upregulated in macrophages upon infection and requires an active Myd88-immune signalling. (A) Expression of *cxcl11aa* in FACS-sorted macrophages (MΦ, *mpeg1:mCherry* positive) and its infection-dependent induction (relative to negative/Mock fraction). (B) *Mm*-(or Mock-) injected larvae (> 100 per replicate per condition) were dissociated at 5 dpi. Induction of *cxcl11aa* does not require the RD1 pathogenicity locus and mutants of the cognate receptor of cxcl11aa (*cxcr3.2^−/−^*) are still able to upregulate *cxcl11aa* at comparable levels to wt. (C-E) RNA was isolated from pools of > 10 whole larvae collected at 4 dpi. 800 CFU of RD1 mutant bacteria versus 100 CFU of wildtype *Mm* were injected to reach a comparable infection level at 4 dpi. Dependency of *cxcl11aa* induction from *myd88*. Expression levels (C), representative burden analysis (D) and representative burden pictures (E) derive from larvae collected at 4 dpi. RNA was isolated from pools of > 10 whole larvae. Each point in (D) represents 1 infected larva from a representative pool. 200 CFU of wildtype *Mm* were injected in *myd88*^+/+^ larvae versus 100 and 200 CFU injected in *myd88*^−/−^ larvae to reach a comparable infection level at 4 dpi. Quantification of total bacterial pixels was obtained using dedicated bacterial pixel count program (71). Scale bar in (E): 200 μm. Statistical significance was analysed by one-way ANOVA with Sidak post-hoc correction on ln(n)-transformed relative induction folds (real time PCRs) or untransformed data (infection burden). Significance (*P*-value) is indicated with: ns, non-significant; *\*P<0.05; **P<0.01; ***P<0.001; ****P<0.0001.* Error bars: mean±s.e.m.

## Discussion

Zebrafish larvae provide unique possibilities for real time visualization of macrophage responses during developmental and disease processes. However, it has remained unknown how the expression profile of larval macrophages compares to the profiles of human M1 and M2 *in vitro* polarized macrophage subsets, which are commonly considered as a reference for pro-inflammatory or anti-inflammatory activation states. Here we used RNAseq analysis of FACS-sorted cell fractions to determine the expression profile of macrophages isolated from *mpeg1* reporter lines, which are widely used for imaging studies in zebrafish due to the highly specific labeling of the macrophage lineage. We demonstrate the unique signature of the *mpeg1* reporter cells by comparison with the RNAseq profiles of neutrophils, marked by the *mpx* reporter, and progenitor lymphocytes, marked by the *lck* reporter. We detected expression of homologs of human M1 as well as M2 markers in the *mpeg1* reporter cells, indicating that zebrafish larval macrophages have the potential to differentiate into both directions. Finally, to demonstrate polarization of macrophages under challenged conditions, we achieved an RNAseq analysis of low numbers of *mpeg1*-positive macrophages infected with a mycobacterial pathogen. The profiling of these infected macrophages revealed downregulation of M2 markers, while M1 markers were upregulated, with strongest induction of a homolog of the human M1 marker CXCL11.

Adult *mpeg1* reporter fish have previously been used to determine the transcriptome of microglia, the brain-resident macrophage population (60). Other fluorescent reporter lines for different immune cell types from the myeloid and lymphoid lineages have recently been used to determine single-cell transcriptomes of cells sorted from hematopoietic organs (spleen and kidney marrow) of adult fish (61; 62; 63). Our study is the first to report on the transcriptome of larval macrophages. We could detect reproducible expression of a total of 13185 genes in *mpeg1*-positive macrophages, among which 744 showed significantly enriched expression compared to the background tissue. These genes were linked to biological processes that are important for macrophage function, such as innate immunity, inflammation, antigen processing and presentation, signal transduction, peptidase activity, chemotaxis, and actin filament organization and polymerization. This RNAseq dataset provides a useful new data mining resource that will facilitate genetic analyses of macrophage-specific genes in zebrafish larval models for development and disease.

A dual-fluorescent reporter line with *mpeg1*-labelled macrophages and *tnfa* as a marker for M1 phenotype has been used to demonstrate that injury and infection can induce M1 polarization of macrophages in zebrafish larvae (16). While the *tnfa* reporter does not show detectable fluorescent gene expression in the absence of wounding or infection stimuli, we could detect a basal level of *tnfa* expression in our RNAseq data of macrophages from unchallenged zebrafish larvae. Furthermore, our RNAseq data set of 744 enriched macrophage markers contains other M1 markers that were reported to be induced by injury in *tnfa*-positive macrophages (*il1*, *il6*), but also contains M2 markers that are expressed at higher levels in *tnfa*-negative macrophages (*tgfb1*, *ccr2*, *cxcr4b*). Single cell sequencing would be required to determine if all macrophages express these M1 and M2 markers at low levels or that distinct macrophage subsets exist already under unchallenged conditions. A comparison with RNAseq profiles of *in vitro* differentiated human M1 and M2 macrophages provided further evidence that the transcriptome of zebrafish larval macrophages displays a mixed M1 and M2 signature (40). Whereas our results do not allow to conclude if two distinct populations of macrophages similar to human M1 and M2 polarized macrophages exist in zebrafish larvae, we identified several specific genes that suggest the presence of these different populations and that could be used to expand the repertoire of zebrafish transgenic reporter lines for investigating macrophage polarization *in vivo* during immune challenge in the zebrafish model.

Fluorescent reporters driven by the *mpeg1* and *mpx* promoters distinguish specifically between macrophages and neutrophils (20; 27). In agreement, we did not detect *mpeg1* gene expression in neutrophils from *mpx* reporter fish. However, we detected low levels of expression of *mpx* and other common neutrophil markers in macrophages from *mpeg1* reporter fish. This indicates that the RNAseq procedure is highly sensitive and suggests that post-translational mechanisms contribute to regulating the specificity of innate immune cell types. We found that approximately half of the genes that show enriched expression in neutrophils also show enriched expression in macrophages. However, an obvious difference between the two innate immune cell types is that genes involved in antigen presentation and processing were detected only in macrophages. Other notable differences were found within the families of proteinases, with macrophages expressing several members of the cathepsin family, while neutrophils express genes from the carboxypeptidase, elastase, and chymase families. The neutrophil RNAseq data reported here have been data mined to investigate the expression of the major classes of drug transporters in zebrafish larvae, providing useful information for optimizing screening approaches for anti-inflammatory drugs (64).

The enriched gene sets of larval macrophages and neutrophils consist for more than 75% of transcripts that are not detected in progenitor lymphocytes isolated from *lck* reporter fish. Similarly, the enriched gene set of *lck*-labelled lymphocytes consists for 85% of transcripts that are not expressed in the myeloid lineage. It is well known that the adaptive immune system in zebrafish larvae is not yet mature and that full immunocompetence, including antibody production, is achieved only by 3-6 weeks of development (65; 66). However, it is not known at which stage of larval development the first interactions between antigen-presenting cells and T-lymphocytes take place. We found that *mpeg1*-labelled macrophages from 5 day old larvae express the major histocompatibility class II gene *mhc2dab*, which is even earlier than the observed expression of a fluorescent *mhc2dab* reporter that labels putative dendritic cells scattered throughout the skin of larvae from 9 days onwards (67). The presence of *mhc2dab* and transcripts of other genes involved in antigen presentation and processing in larval macrophages suggest that communication with T-lymphocytes could take place already at stages where zebrafish larvae are generally believed to rely exclusively on innate immunity. In support of this hypothesis, we found that larval lymphocytes express the Cd4 marker for helper T-cells and the co-stimulatory receptor Cd28, which are required for macrophage activation. Furthermore, the expression of perforin genes is indicative of the development of cytotoxic T-cells. However, since there was no detectable expression of Cd8, it is unlikely that cytotoxic T-cells are already functional in 5 day old larvae.

To investigate how larval macrophages respond to an intracellular infection with mycobacteria, we determined the expression profile of *Mm*-infected *mpeg1* reporter cells. This RNAseq analysis was challenging due to the low numbers of infected cells that could be obtained by FACS sorting. Infected macrophages have a lifespan of less than 5 hours (68), which likely is an important contributing factor to the difficulty of isolating *Mm*-infected cells. To deal with the complication of low cell numbers, we adapted the RNAseq protocol and determined the transcriptomes of triplicate pools of 20 infected or non-infected cells from *M. marinum* infected larvae. By limiting the number of cells per sample to 20 we could avoid interference of DNA reads in the RNAseq data, while still keeping sufficient biological variation to answer the question if *Mm* infection causes a general shift in macrophage polarization. While different types of macrophage polarization have been reported for *in vitro* cultured macrophages infected with mycobacteria (15), it is not understood how these pathogens affect macrophage polarization during different stages of tuberculosis disease *in vivo*. Our RNAseq analysis was performed at a stage where infected macrophages have aggregated into inflammatory infection foci, which are regarded as the earliest developmental stages of tuberculous granulomas (69). We observed that homologs of M2-enriched transcripts of human cells were preferentially down-regulated in *M. marinum-*infected zebrafish macrophages, whereas several M1-enriched transcripts were highly upregulated. Therefore, although no clear polarization was observed, our analysis suggests that macrophages shift towards M1 phenotype in *Mm*-infected zebrafish, which are used to model tuberculosis. Our results show an important modification of the macrophage transcriptome upon mycobacterial infection and unravel several targets that can be studied to better understand the molecular mechanisms involved in the host-pathogen interaction.

An important question is whether part of the observed expression changes in *Mm*-infected macrophages might be triggered by bacterial virulence factors or that all changes represent a general host defense response that is mounted against pathogenic as well as non-pathogenic mycobacteria. Irrespective of the answer to this question, it can be argued that some of the induced genes benefit the pathogen rather than the host. For example, we detected induced expression in *Mm*-infected macrophages of genes (*nsfb*, *acap1*) that promote the survival of bacteria in phagosomes (55; 56). A gene that is induced strongly and reproducibly among all replicates, *cxcl11aa*, could have both host-beneficial and host-detrimental effects. This gene, which is a homolog of the human M1 marker *CXCL11*, encodes a chemokine that mediates macrophage recruitment to infection foci through interaction with chemokine receptor Cxcr3.2, the zebrafish counterpart of human CXCR3 (39). While a certain level of macrophage recruitment during *Mm* infection is necessary to restrict infection (70), *Mm* bacteria also take advantage of the arrival of new macrophages at infection foci as this promotes spreading of the infection (37). In line with these considerations, we have previously found that deficiency in the receptor for Cxcl11aa, Cxcr3.2, limits the expansion of *Mm* in granulomas (39). A similar phenotype has been found upon depletion of Mmp9, another host factor required for macrophage recruitment (38). Therefore, high and sustained induction of *cxcl11aa* is likely to have an adverse effect on the control of *Mm* infection by the zebrafish host. On the other hand, the robust induction of this M1 polarization marker makes the *cxcl11aa* gene a prime candidate to expand the collection of zebrafish reporter lines for studying macrophage activation.

In conclusion, the transcriptome analyses reported here present a unique and detailed genetic profile of zebrafish larval immune cells, thereby providing a valuable resource that can be data mined to verify the expression of specific genes in the profiled cell types or to identify novel genes of interest and potential cell-specific markers. In future work single cell RNA sequencing technology will be useful to interrogate the heterogeneity in expression profiles of resting and activated macrophages.

## Data Availability Statement

The sequencing data for infected samples have been submitted to the NCBI Gene Expression Omnibus (GEO; http://www.ncbi.nlm.nih.gov/geo/) under accession number GSE68920. The sequencing data for uninfected samples were made previously available under accession number GSE78954.

## Acknowledgements

The authors thank Michiel van der Vaart for critical reading of the manuscript, Graham Lieschke (Monash University), Georges Lutfalla (University of Montpellier), Steve Renshaw (University of Sheffield) and David Langenau (Harvard Stem Cell Institute) for zebrafish reporter lines, Guido de Roo and Sabrina Veld for support with FACS sorting, and the fish facility team members for zebrafish care. This work was supported by the Marie Curie Initial Training Network FishForPharma (PITN-GA-2011-289209) and the project ZF-HEALTH (HEALTH-F4-2010-242048) funded by the European Commission 7th Framework Programme, and by the SmartMix programme of the Netherlands Ministry of Economic Affairs and the Ministry of Education, Culture, and Science. Additionally, Z.K. was supported by the Higher Education Commission of Pakistan and A.Z. was supported by a Horizon grant of the Netherlands Genomics Initiative.

## Author Contributions

JR contributed to conception and design of the study, performed and analyzed experiments, and wrote and edited the manuscript. VT contributed to conception and design of the study, performed and analyzed experiments, and wrote and edited a section of the manuscript. AZ contributed to conception and design of the study and performed experiments. ZK contributed to conception and design of the study and performed experiments. HJJ performed and analyzed experiments. AHM contributed to conception and design of the study, wrote and edited the manuscript, and acquired funding. All authors contributed to manuscript revision, read and approved the submitted version.

## Conflict of Interest Statement

Author HJJ was employed by company ZF-screens B.V. (Leiden, The Netherlands). All other authors declare no competing interests.

## Supplementary Material

Figure S1. Correlation between RNA-sequencing samples. (A-C) HeatMap of Pearson correlation coefficient of 5 and 6 dpf *mpeg1:gal4; UAS-Kaede* and *mpeg1:mCherry* positive and negative samples (A), 5dpf *mpx:gfp* and *lck:gfp* positive and negative samples (B) and 20 cell samples of *mpeg1:gfp* positive cells infected or not with *M. marinum* GFP (C).

Figure S2. Principal Component Analysis of RNA-sequencing samples. (A-C) Principal Component Analysis of 5 and 6 dpf *mpeg1:gal4; UAS-Kaede* and *mpeg1:mCherry* positive and negative samples (A), 5dpf *mpx:gfp* and *lck:gfp* positive and negative samples (B), and 20 cell samples of *mpeg1:gfp* positive cells infected or not with *M. marinum* GFP (C).

Figure S3. Molecular function associated to the zebrafish core macrophage expression dataset. Network visualization of GO analysis enrichment (molecular function category) of genes from the core macrophage expression data set using BiNGO and EnrichmentMap. Node size corresponds to the number of genes associated to the enriched GO term and edge size to the similarity coefficient between two nodes.

Table S1. Additional gene annotation. List of manually annotated genes added to the Ensembl annotation version 79 from the genome version Zv9.

Table S2. FPKM table of macrophage, neutrophil and lymphoid cell population. FPKM for each positive and negative samples were computed by using the longest transcript for each gene. Presented results are average between each replicate.

Table S3. Summary of gene expression and differential expression analysis. Level of gene expression was distributed based on average FPKM values among non-expressed (FPKM<0.3), moderately expressed (0.3≤FPKM<100) and highly expressed (FPKM≥100) for the positive samples and an average of all the negative samples present in this analysis. Fold Change (FC) between positive and negative samples were computed using DESeq as described in the material and methods.

Table S4. Zebrafish core macrophage expression dataset. The 744 genes enriched (*P*-adj < 0.05) in at least two of the three *mpeg1* positive cell populations compared with their respective negative cell background represent the zebrafish core macrophage expression data set. Associated gene names and descriptions in bold are additional manual annotations to the Ensembl annotation. For each gene, one relevant term from the Biological Process GO is presented.

Table S5. Gene expression dataset in neutrophils. List of genes enriched (FC > 0 and *P*-adj < 0.05) in *mpx:gfp* positive neutrophils compared with fluorescent negative cells. Associated name and description in bold are additional annotations to the Ensembl annotation. For each gene, one relevant term from the Biological Process GO is presented.

Table S6. Gene expression dataset in lymphoid cells. List of genes enriched (FC > 0 and *P*-adj < 0.05) in *lck:gfp* positive lymphoid cells compared with fluorescent negative cells. Associated name and description in bold are additional annotations to the Ensembl annotation. For each gene, one relevant term from the Biological Process GO is presented.

Table S7. List of the human homologs from zebrafish genes. List of the human homologs from the genes enriched in the zebrafish core macrophage expression dataset.

Table S8. Differential expression analysis between *Mycobacterium marinum* infected and uninfected zebrafish macrophages. List of genes upregulated (FC > 0 and *P*-adj < 0.05) and downregulated (FC < 0 and *P*-adj < 0.05) in *mpeg1:mCherry* and *Mm-GFP* double positive macrophages compared with *mpeg1:mCherry* only positive macrophages. Associated name and description in bold are additional annotations to the Ensembl annotation.

