## Supplementary figures and images for "RNAseq profiling of leukocyte populations in zebrafish larvae reveals a *cxcl11* chemokine gene as a marker of macrophage polarization during mycobacterial infection"

### Supplemental Figure 1

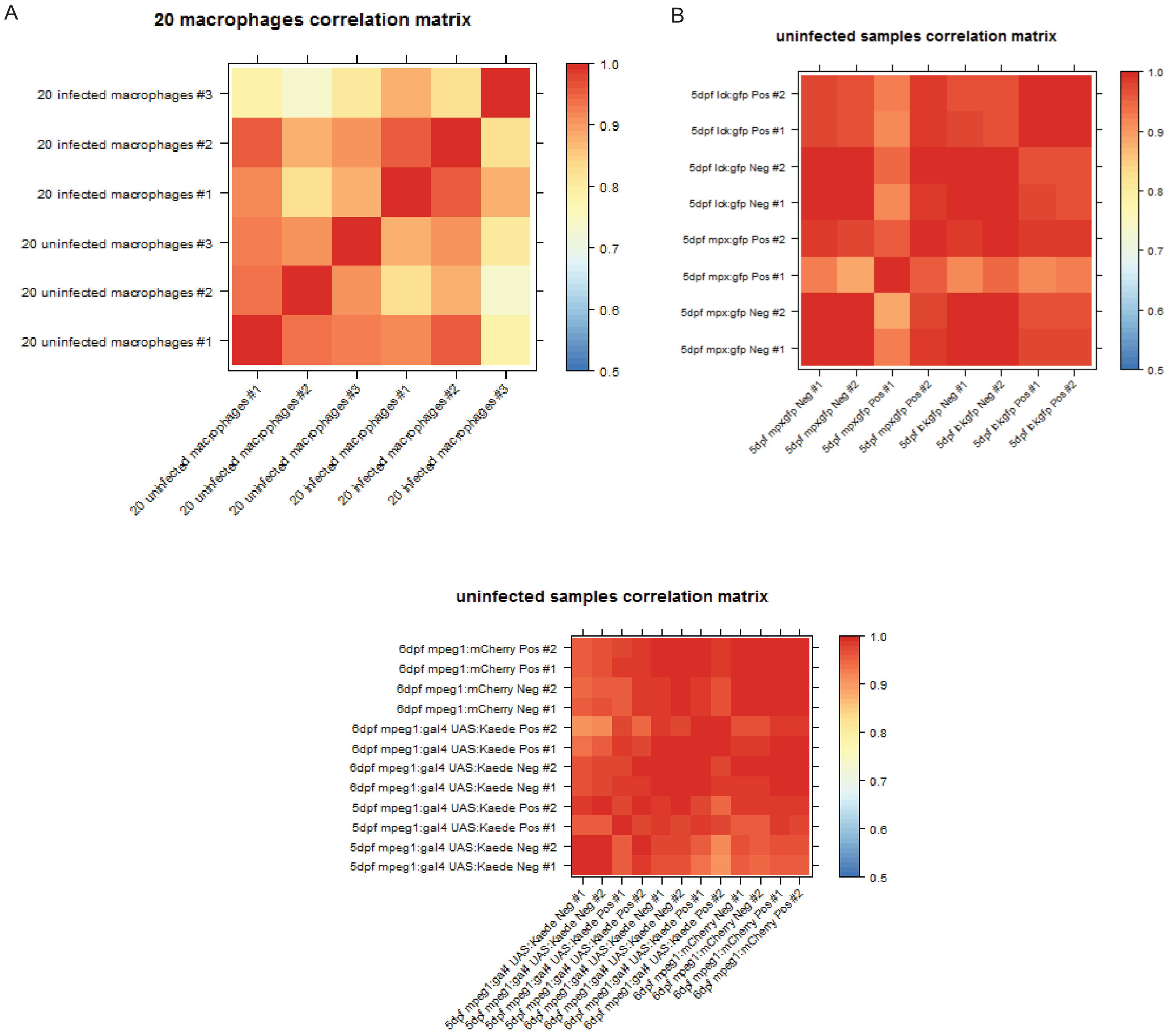


Rougeot_et_al.: Supplementary figure 1

### Supplemental Figure 2

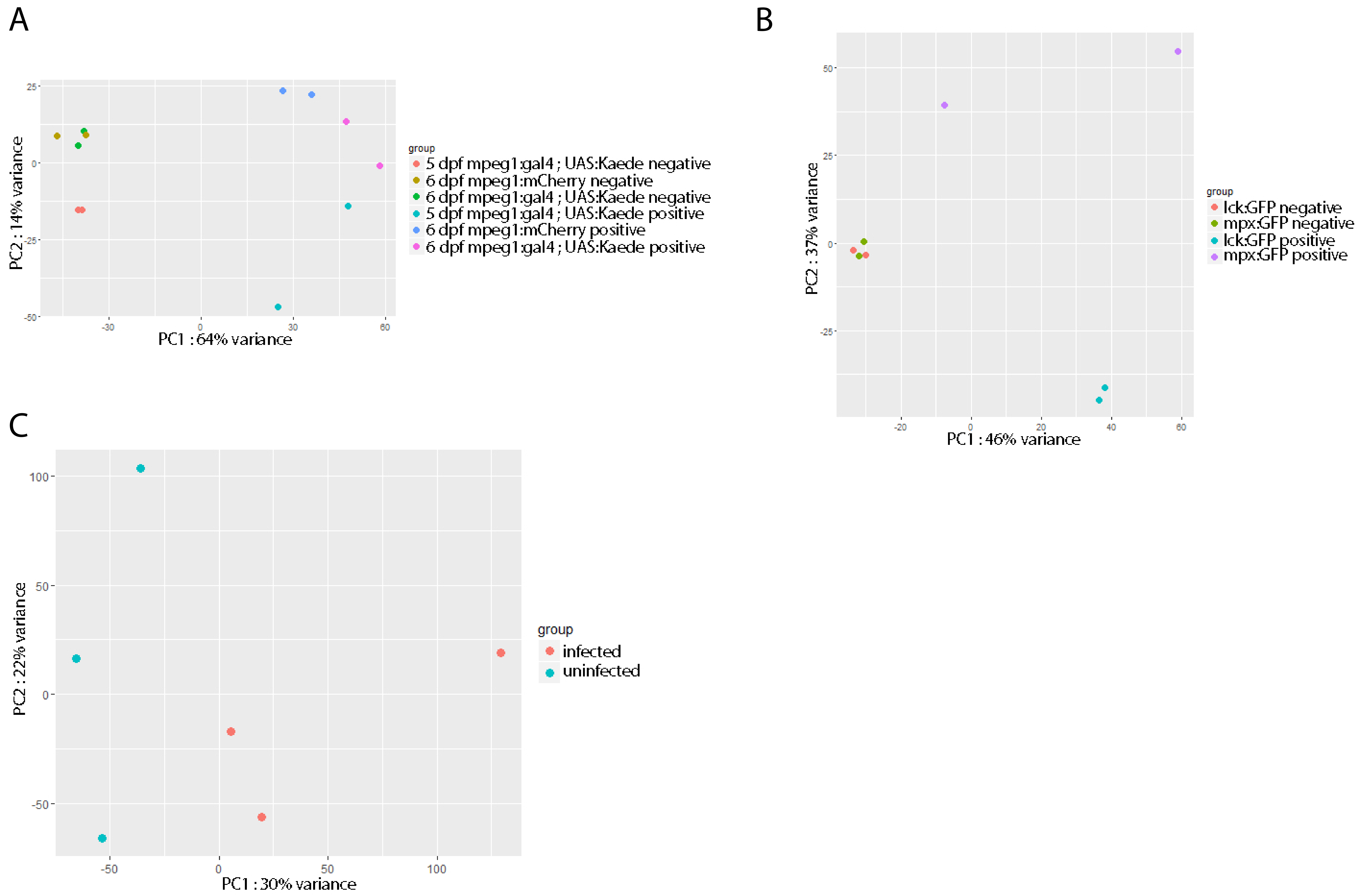


Rougeot_et_al.: Supplementary figure 2

### Supplemental Figure 3

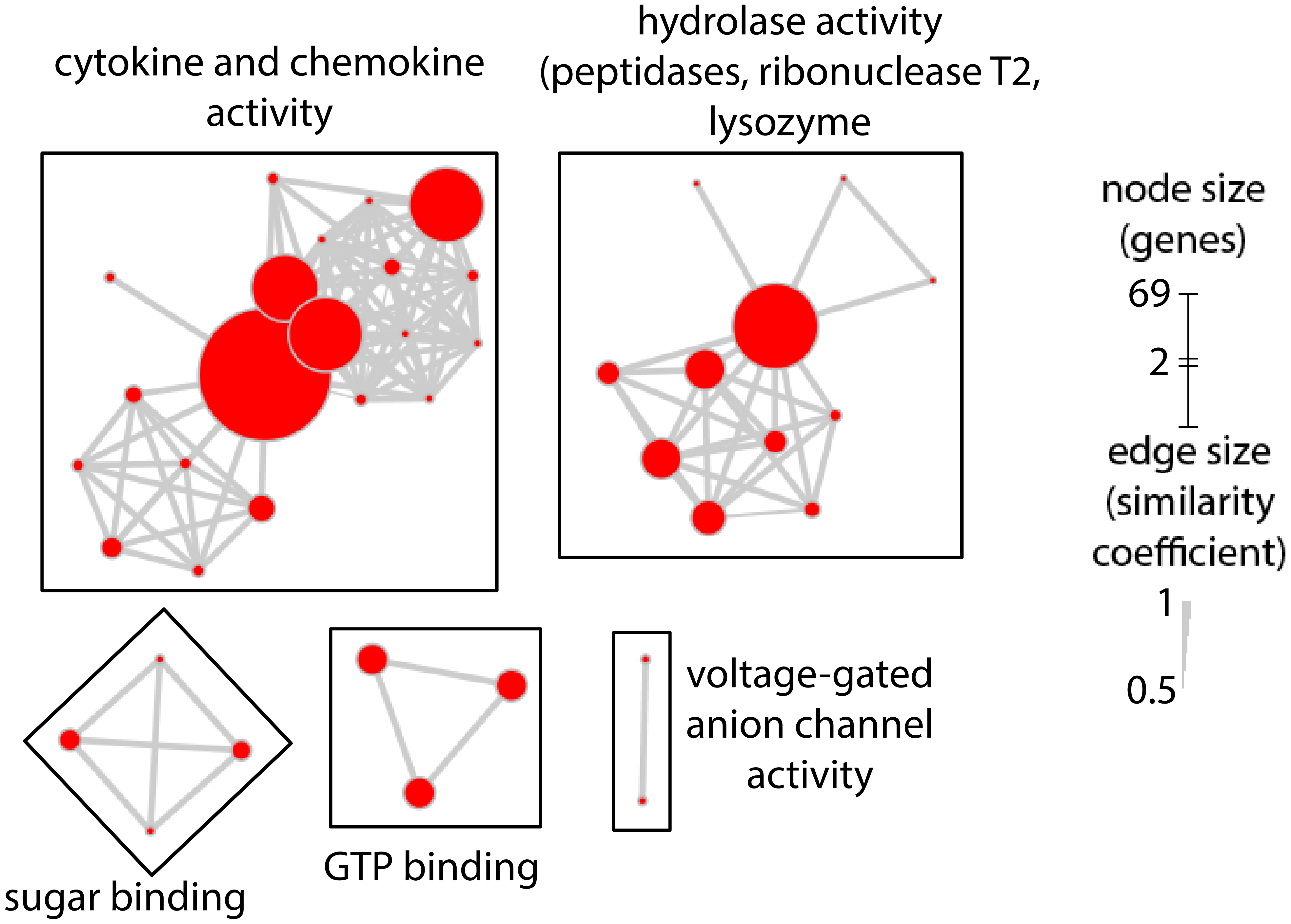


Rougeot_et_al.: Supplementary figure 3
